# Genetic alteration of ferredoxin NADP^+^-reductase or cysteine desulfurase in piperaquine-resistant *Plasmodium berghei* restores susceptibility to lumefantrine and abolishes the impact of probenecid, verapamil, and cyproheptadine

**DOI:** 10.1101/833145

**Authors:** Fagdéba David Bara, Loise Ndung’u, Noah Machuki Onchieku, Beatrice Irungu, Simplice Damintoti Karou, Francis Kimani, Damaris Matoke-Muhia, Peter Mwitari, Gabriel Magoma, Alexis Nzila, Daniel Kiboi

## Abstract

Chemotherapy remains central in the control of malaria; however, resistance has consistently thwarted these efforts. Currently, lumefantrine (LM), and piperaquine (PQ) drugs, are essential components in the mainstay artemisinin-based therapies used for the treatment of malaria globally. Using LM and PQ-resistant *Plasmodium berghei*, we measured the effect of known chemosensitizers: probenecid, verapamil, or cyproheptadine on the activity of LM or PQ. Using PlasmoGEM vectors, we then evaluated the impact of deleting cysteine desulfurase (SufS) or over-expressing Ferredoxin NADP+ reductase (FNR), genes that mediate drug action. Our data showed that, only cyproheptadine at 5mgkg^−1^ restored LM activity by above 65% against the LM-resistant parasites (LM^R^) but failed to reinstate PQ activity against the PQ-resistant parasites (PQ^R^). Whereas the PQ^R^ had lost significant susceptibility to LM, the three chemosensitizers; cyproheptadine, probenecid, and verapamil, restored LM potency against the PQ^R^ by above 70%, 60%, and 55% respectively. We thus focused on LM resistance in PQ^R^. Deletion of the SufS or overexpression of the FNR genes in the PQ^R^ abolished the impact of the chemosensitizers on the LM activity, and restored the susceptibility of the PQ^R^ parasites to LM. Taken together, we demonstrate the association between SufS or FNR genes with the action of LM and chemosensitizers in PQ^R^ parasites. There is, however, need to interrogate the impact of the chemosensitizers and the role of SufS or FNR genes on LM action in the human malaria parasite, *Plasmodium falciparum*.

## INTRODUCTION

Malaria affects more than one billion people worldwide ^1^. Presently, five malaria parasite species infect humans. *Plasmodium falciparum* remains a significant contributor to the global disease burden, with an estimated 228 million cases and 405 000 deaths annually ^1^. An enormous proportion of this burden affects children under five years of age and pregnant women. In Africa, over 70% of the population is still at risk of infection by the malaria parasites ^1,2^. To date, the use of drugs is central to the control and management of malaria. However, this approach is hampered by the ability of the parasite to rapidly develop resistance against antimalarial drugs. Currently, the artemisinin-based combination therapies (ACTs) are the mainstay drugs for the treatment and management of uncomplicated *P. falciparum* malaria. The ACTs comprise a short-acting artemisinin derivative and a long-acting partner drug ^2^. The long-acting partner drugs reduce the remaining parasite biomass after artemisinin clearance and simultaneously protect against reinfection, especially in high transmission settings ^3^. Thus, the long-acting drug components within the ACTs are of primary importance in the control of subsequent malaria infection in sub-Saharan Africa.

In many African countries, the long-acting antimalarial drugs: lumefantrine (LM) and piperaquine (PQ) are essential components in the mainstay ACTs; Coartem™ and Artekin™, respectively ^4^. Unfortunately, the high transmission of malaria in endemic regions coupled with the long half-life portend intense selection pressure ^5^, a recipe for the rapid emergence of LM and PQ resistant parasites. It is therefore imperative to understand how parasites may evade LM and PQ action. Polymorphisms in two genes, primarily mediate resistance to the 4-aminoquinolines and chemically related drugs such as PQ. The *chloroquine resistance transporter (crt)*, the fundamental determinant of CQ resistance in *P. falciparum*, which can carry Lys76Thr mutation ^6^ and *multidrug-resistant 1 (mdr1)* gene that encodes a P-glycoprotein homolog 1 (*Pgh-1*) which modulate resistance to CQ and other quinolines drugs such as AQ and PQ ^7,8^. In recent studies, Cys101Phe mutation in *chloroquine resistance transporter* (*crt*) conferred resistance to both PQ and CQ in *P. falciparum in vitro* ^9^. Also, using field isolates, a nonsynonymous mutation, Glu415Gly in the exonuclease (PF3D7_1362500), and amplification of Plasmepsin II and III, proteases involved in the heme degradation within the digestive vacuole are linked with PQ resistance ^10^. LM, on the other hand, is an aryl alcohol antimalarial drug, chemically related to mefloquine and quinine. Although, like the 4-aminoquinoline, LM is predicted to inhibit heme detoxification, several studies have associated reciprocal resistance between CQ and LM ^11^, suggesting potentially different mechanisms of resistance and action. To date, many questions regarding how LM exerts its antimalarial action and how the parasites might evolve to evade such action remain unanswered.

One archetypical method of studying the mechanisms by which parasites evade drug action and simultaneously generate new drugs is the use of a chemosensitizer (a drug with the ability to enhance/restore activity) in association with antimalarial drugs for which the parasite is resistant ^12^. This approach has been utilized previously to generate prophylactic drugs against malaria parasites ^13,14^. Although many chemosensitizers have no intrinsic antimalarial potency, an antihistamine cyproheptadine can reverse resistance and kill asexual blood-stage malaria parasite^15^. Chemosensitizers reverse resistance mechanisms by either enhancing drug uptake or inhibiting the efflux of drugs from the target site ^16–18^. Three chemosensitizers have been extensively used to study the resistance mechanisms and design next-generation drugs in both malaria and cancer ^19–22^. In malaria, verapamil, a calcium (Ca^+2^) channel blocker, partially reverses CQ resistance in *P. falciparum in vitro* and in clinical isolates via increasing net CQ within the infected erythrocyte parasites ^23^. Probenecid, anion transporter inhibitor, reverses methotrexate resistance in cancer ^24^, CQ, and antifolates resistance in *P. falciparum* ^14^. Cyproheptadine and a related first-generation antihistamine; chlorpheniramine restore CQ sensitivity in CQ resistant *P. yoelii* and *P. falciparum in vitro* ^15,25^.

Using a model malaria parasite, *Plasmodium berghei* ANKA, we have previously selected PQ- and LM-resistant parasites^26^. Further analysis of the PQ^R^ parasites showed that the parasite had lost significant susceptibility to LM, amodiaquine (AQ), primaquine (PMQ), and dihydroartemisinin (DHA)^26^. On the other hand, although LM^R^ parasites retained sensitivity to PQ, the parasites lost significant susceptibility to DHA, PMQ, and CQ ^26,27^. Also, the resistant parasites acquired significant growth fitness cost probably associated with the changes within the genomes that affect the essential genes necessary for the growth of the asexual blood-stage parasites ^28^. In the current study, we hypothesized that: i) a combination of probenecid, verapamil, or cyproheptadine with PQ might restore the efficacy of PQ against the PQ^R^ parasites; ii) a combination of probenecid, verapamil, or cyproheptadine with LM might restore the efficacy of LM against both PQ^R^ and LM^R^ parasites; iii) PQ^R^ and LM^R^ parasites are accompanied by differential mRNA expression of a panel of plausible genes that mediate drug action and transport quinoline and aryl alcohol drugs. These are; *crt* (PBANKA_1219500), *mdr1* (PBANKA_1237800), *multidrug resistance-associated protein 2 (mrp2)* (PBANKA_1443800), *pantothenate kinase (pank)* (PBANKA_1022600), *ferredoxin NADP^+^ -reductase (fnr)* (PBANKA_1122100), *cysteine desulfurase (SufS)* (PBANKA_0614300), *Plasmepsin IV (pmiv)* (PBANKA_1034400), *Plasmepsin IX (pmix)* (PBANKA_1014500), *Plasmepsin X (pmx) or phosphatidylinositol 3-kinase (pi3k)* (PBANKA_1114900); iv) deletion or overexpression of selected plausible genes would restore parasite susceptibility to PQ or LM. Here, we provide insights into the mechanisms of LM resistance and reversal in *Plasmodium berghei*, which are critical for future rational drug design.

## RESULTS

### Cyproheptadine selectively restores LM activity against lumefantrine and piperaquine resistant parasites

To select the appropriate concentrations for the drug combination studies, we first separately conducted cyproheptadine, LM or PQ activity assays against the resistant (PQ^R^ and LM^R^) and wildtype (WT) parasites. Since the LD_50_ of cyproheptadine is 123mg/kg in mice^29^, we used 30mg/kg as the highest concentration. Surprisingly, 30mg/kg of cyproheptadine killed 90% of the LM^R^ (Fig 1A) and 40% of the PQ^R^ parasites (Fig 1B). However, at 5mg/kg, cyproheptadine had no significant activity against either the WT (4.4%), LM^R^ (0.6%) or PQ^R^ parasites (0.7%) (Fig 1A and Fig 1B). Based on these cyproheptadine data, a concentration of 5mg/kg or 2.5mg/kg was selected for subsequent drug combination assays. To select the appropriate concentrations of LM or PQ for use with cyproheptadine, we measured the activity profiles of LM against LM^R^ or WT and PQ against PQ^R^ parasites. As expected, 50mg/kg of LM yielded a marginal effect (1.66% parasite killing) on the LM^R^ parasites. Against the WT, 0.375mg/kg of LM showed insignificant parasite killing (17%) compared to 5mg/kg, which yielded 100% parasite killing (Fig 1A, Table 1). On the other hand, 75mg/kg of PQ killed a meager 10.1% of the PQ^R^ parasites as expected (Fig 1B). Based on these LM and PQ activity results, we selected 100mg/kg, 50mg/kg or 25mg/kg of LM for LM^R^ parasites assays, 0.375mg/kg of LM for WT parasites assays, and 75mg/kg of PQ for PQ^R^ parasites assays, for subsequent drug combination experiments.

**Fig 1:**
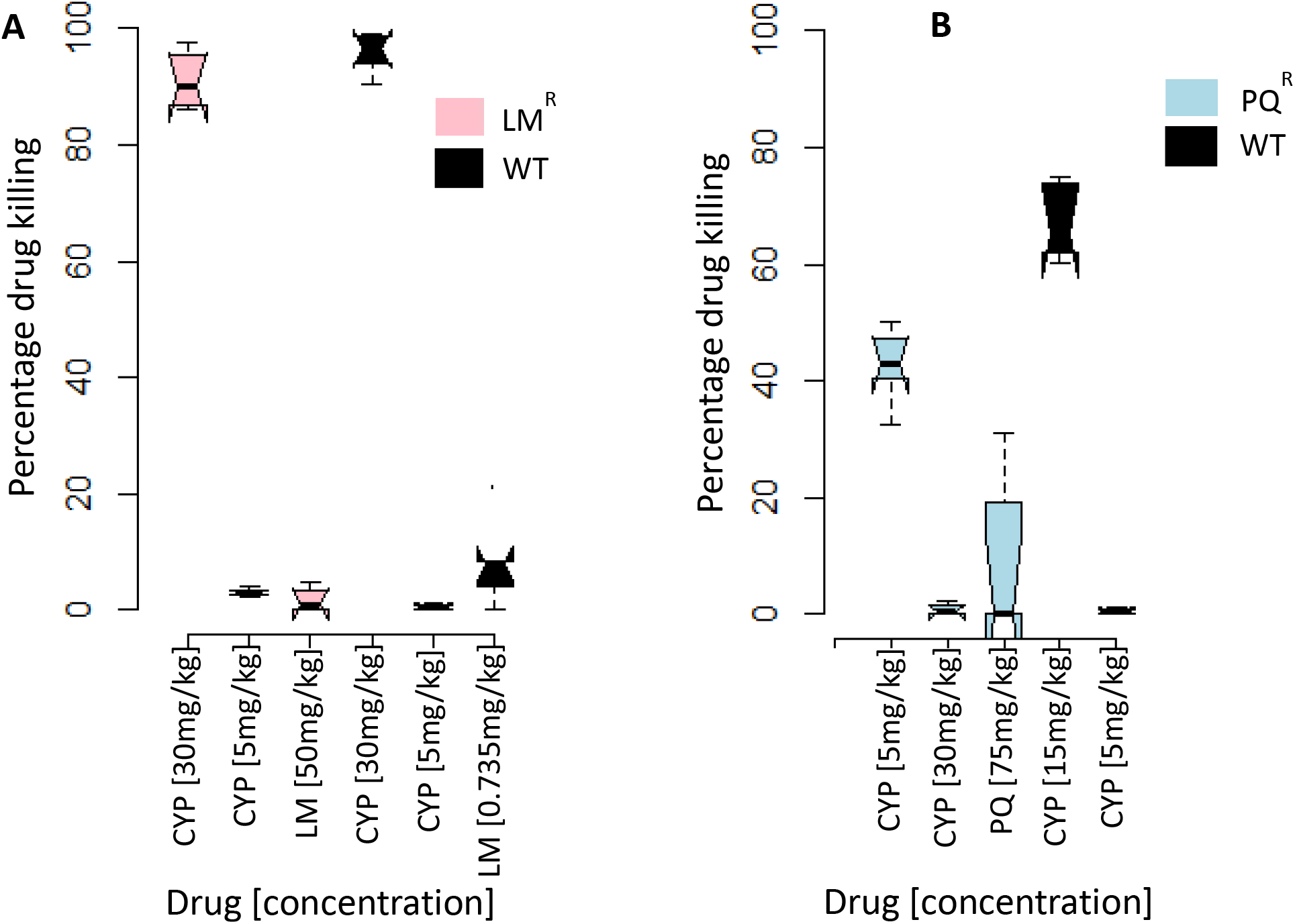
Activity profile of cyproheptadine (CYP), Lumefantrine (LM) and piperaquine (PQ) as assessed in the 4-Day Suppressive Test. **(A)** CYP and LM against the lumefantrine resistant (LM^R^) and wildtype parasites (WT). **(B)** CYP and PQ against the piperaquine resistant (PQ^R^) and wildtype (WT) parasites. The results are presented as percentage (%) drug killing of parasites in at least five mice per dose.

**Table 1:** Effects of cyproheptadine (Cyp) or lumefantrine (LM) alone or in combination against the Wildtype (WT) *Plasmodium berghei* ANKA as assessed in the 4-Day-Test. Results presented as percentage (%) parasite killing ± SD (standard deviation) of five mice per dose

| Drug | Dose (mg/kg) | % chemo-suppression ± SE |
| --- | --- | --- |
| Cyp | 5 | 4.44 ± 1.65 |
| Cyp | 2.5 | 5.32 ± 4.12 |
| LM | 0.375 | 17.12 ± 4.12 |
| LM | 5 | 100.0 |
| LM + Cyp | 0.375 + 5 | 20.77 ± 5.25 |
| LM + Cyp | 0.375 + 2.5 | 5.93 ± 2.57 |

We first measured the impact of cyproheptadine on LM activity against the LM^R^ parasites. Here, a combination of cyproheptadine [5mg/kg] and LM [100mg/kg] significantly restored the LM potency from 23% to 67% (p<0.001) with a surprisingly high standard deviation (Fig 2A). A lower cyproheptadine concentration [2.5mg/kg] and LM [100mg/kg] yielded 91% parasite killing with low standard deviation (Fig 2A). Next, we hypothesized that the restoration of LM potency by cyproheptadine was dependent on the concentration of LM or cyproheptadine. To dissect this hypothesis, we evaluated cyproheptadine at 2.5 and 5mg/kg with 25 or 50mg/kg of LM. Cyproheptadine [2.5mg/kg] and LM [25mg/kg] yielded 67%, cyproheptadine [2.5mg/kg] and LM [50mg/kg] yielded 60.6%, cyproheptadine [5mg/kg] and LM [25mg/kg] yielded 61% while cyproheptadine [5mg/kg] and LM [50mg/kg] yielded 68% of parasite clearance. Statistical analysis revealed that the % parasite killing for the different combinations was not significantly different (p<0.08) (Fig 2B), signifying that the restoration of LM activity by cyproheptadine was dose-independent. These data also reaffirmed 2.5mg/kg of cyproheptadine was sufficient low concentration to restore at least two-thirds of LM efficacy against the LM^R^ parasites without obvious reciprocal drug interaction. We next reasoned that the reversal of LM resistance was selective only in the resistant parasites. To evaluate the selectivity of cyproheptadine on the resistant parasites, 2.5 and 5mg/kg of cyproheptadine was combined with 0.375mg/kg of LM against the wildtype parasites. Cyproheptadine [2.5mg/kg] and LM [0.375mg/kg] killed a paltry 6% of the parasites while cyproheptadine [5mg/kg] and LM [0.375mg/kg] yielded 20% compared to 17% of LM [0.375mg/kg] alone (Table 2). Neither of the cyproheptadine concentration increased the susceptibility of the wildtype parasites to LM (p<0.09), meaning that the potentiation of LM activity was selective only in the resistant parasites.

**Fig 2:**
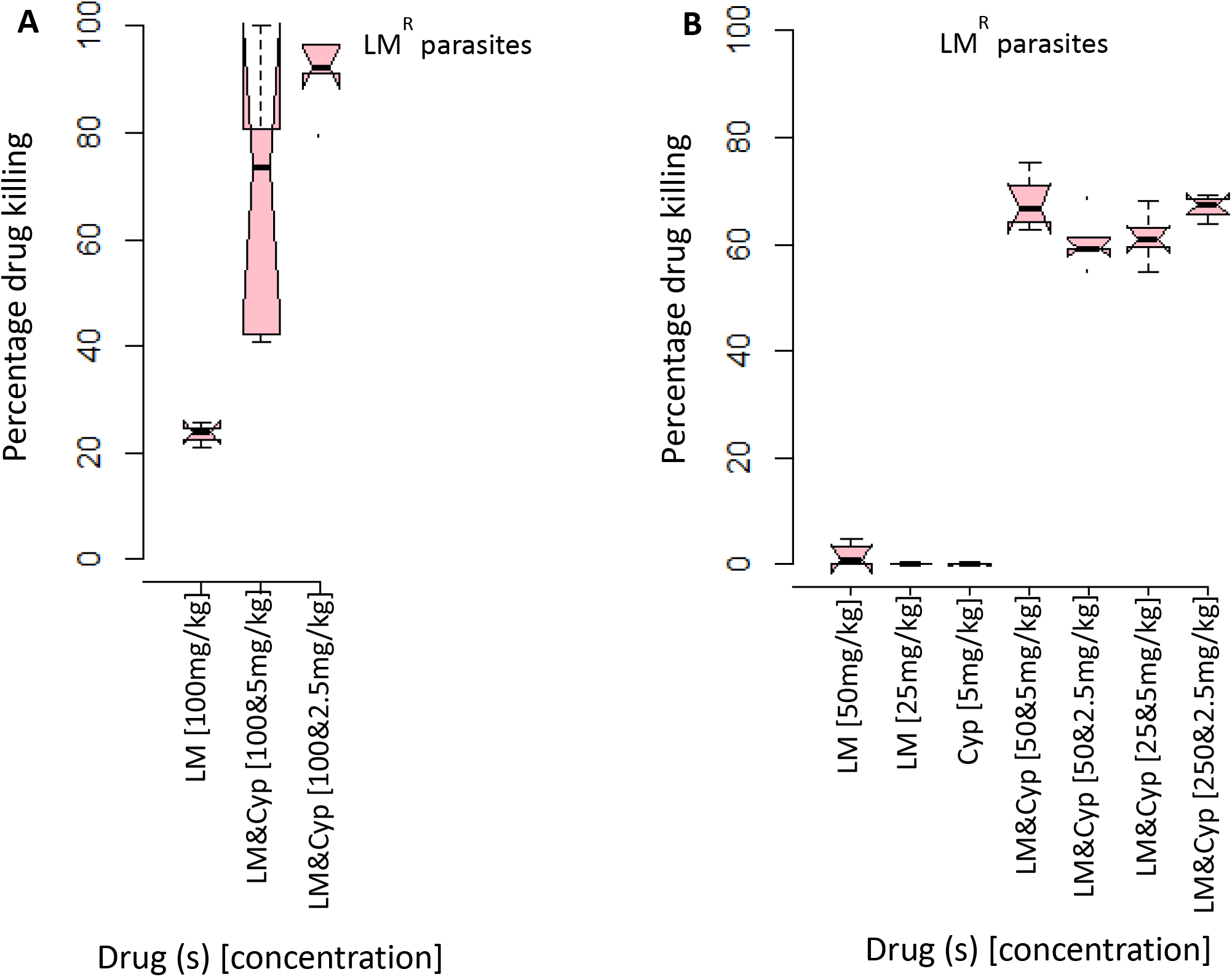
Effect of cyproheptadine (Cyp) on lumefantrine (LM) against the lumefantrine resistant (LM^R^) parasites as assessed in the 4-Day Suppressive Test. **(A)** Impact of varying concentration of cyproheptadine on LM activity against LM^R^ parasites **(B)** Cyproheptadine restores LM potency against LM^R^ parasites independent of synergy and additive effects. The results are presented as percentage (%) drug killing of parasites in at least five mice per dose.

**Table 2:** Effect of verapamil (Ver) and probenecid (Prob) alone against piperaquine resistant (PQ^R^) and lumefantrine resistant (LM^R^) *Plasmodium berghei* ANKA parasites as assessed in the 4-Day Test. Results presented as percentage (%) parasite killing ± SD (Standard Deviation) of five mice per dose.

| Reversing agent | Dose (mg/kg) | % chemo-suppression ± SE<br>(LM<sup>R</sup>) | % chemo-suppression ± SE<br>(PQ<sup>R</sup>) |
| --- | --- | --- | --- |
| Probenecid (Prob) | 400 | 8.44 ± 3.07 | 8.03 ± 1.86 |
| | 200 | 0.00 | 4.06 ± 2.75 |
| Verapamil (Ver) | 50 | 0.00 | 0.81 ± 1.31 |
| | 25 | 0.00 | 0.78 ± 1.56 |

Our next combination assays measured the impact of cyproheptadine on PQ or LM activity against the PQ^R^ parasites. On average, 75mg/kg of PQ yielded 8% parasite killing, a combination of cyproheptadine [5mg/kg] and PQ [75mg/kg] yielded 14% while cyproheptadine [2.5mg/kg] and PQ [75mg/kg] showed 8% parasite killing. We thus concluded that cyproheptadine failed to restore the activity of PQ against PQ^R^ parasites (Table 3).

**Table 3:**
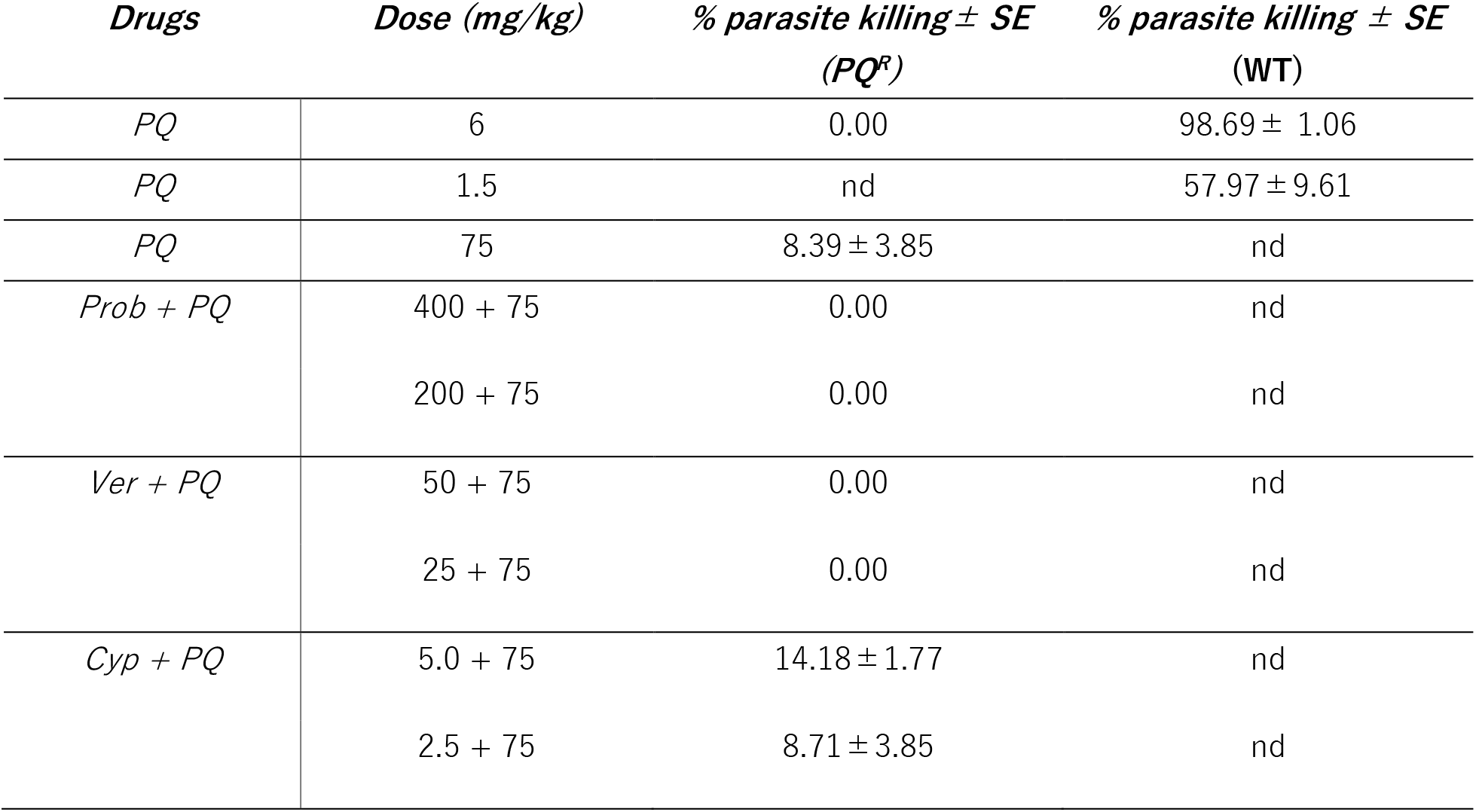
Activity profile of piperaquine (PQ) alone against the wild type (WT) and PQ-resistant (PQ^R^) *Plasmodium berghei* ANKA or PQ in combination with probenecid (Prob), verapamil (Ver), or cyproheptadine (Cyp) against the PQ^R^ parasites assessed in the 4-day Test. Results presented as percentage (%) parasite killing ± SD (standard deviation) of five mice per dose

| *Drugs* | *Dose (mg/kg)* | *% parasite killing ± SE*<br>*(PQ<sup>R</sup>)* | *% parasite killing ± SE*<br>*(WT)* |
| --- | --- | --- | --- |
| *PQ* | 6 | 0.00 | 98.69 ± 1.06 |
| *PQ* | 1.5 | nd | 57.97 ± 9.61 |
| *PQ* | 75 | 8.39 ± 3.85 | nd |
| *Prob + PQ* | 400 + 75 | 0.00 | nd |
|  | 200 + 75 | 0.00 | nd |
| *Ver + PQ* | 50 + 75 | 0.00 | nd |
|  | 25 + 75 | 0.00 | nd |
| *Cyp + PQ* | 5.0 + 75 | 14.18 ± 1.77 | nd |
| | 2.5 + 75 | 8.71 ± 3.85 | nd |

Our previous studies have shown that a significant loss of LM activity accompanies the selection of PQ resistance^26^. We, therefore, investigated whether cyproheptadine might restore LM activity against the PQ^R^ parasites. As expected, 50mg/kg of LM alone had no measurable antimalarial activity (0%) against the PQ^R^ parasites (Fig 3A). Surprisingly, LM activity was re-established by 5mg/kg of cyproheptadine to above 70% (p<0.001) (Fig 3A). The overall results reveal that cyproheptadine selectively reverses LM resistance in LM^R^ parasites but does not restore PQ activity against PQ^R^ parasites. However, it selectively restores LM potency against PQ^R^ parasites.

**Fig 3:**
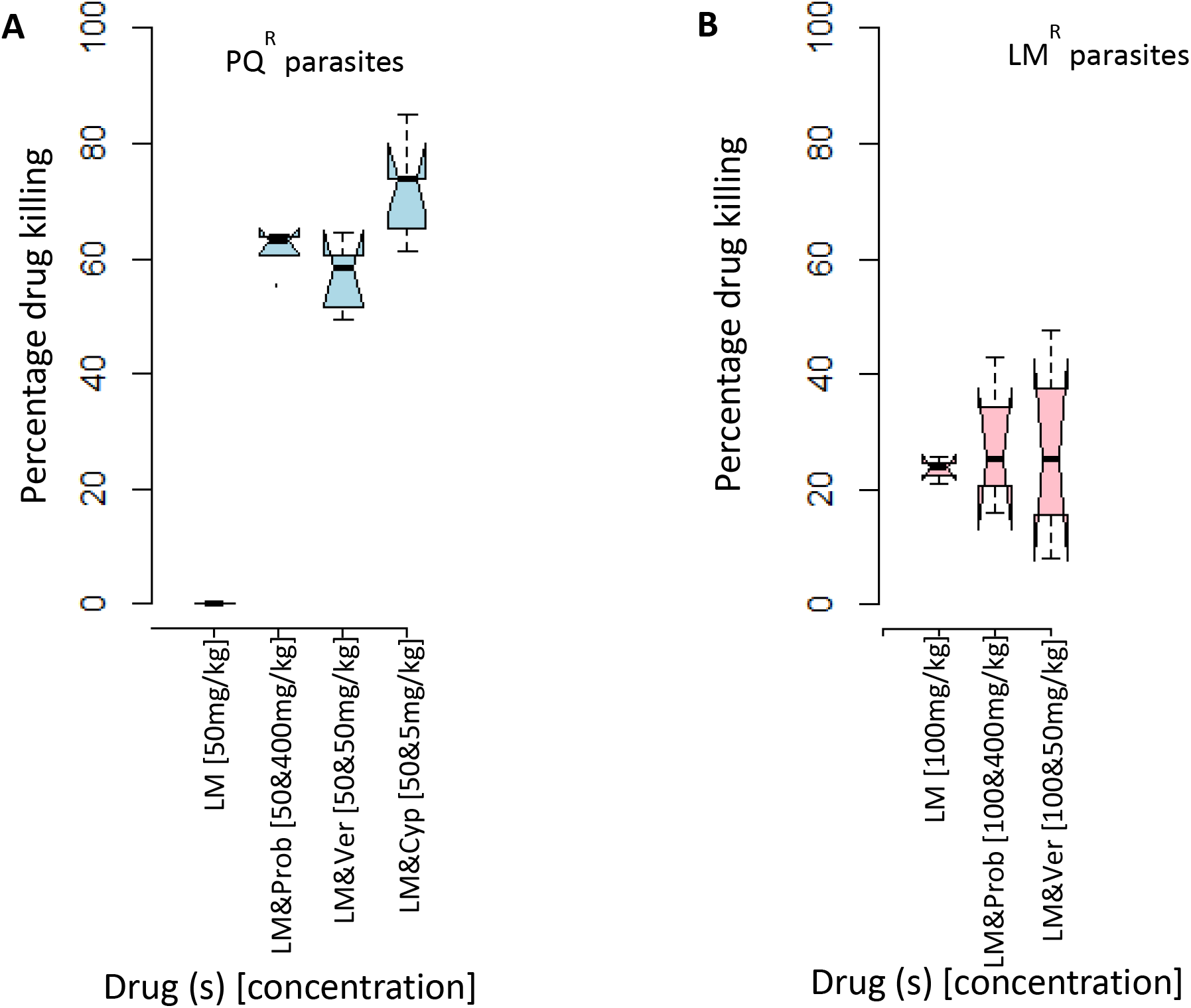
Impact of probenecid (Prob), Verapamil (Ver), or Cyproheptadine (Cyp) on lumefantrine (LM) activity as assessed in the 4-Day Suppressive Test. **(A)** Probenecid, verapamil and cyproheptadine restores LM potency against the piperaquine resistant (PQ^R^) parasites **(B)** Probenecid or verapamil fails to restore LM potency against lumefantrine resistant (LM^R^) parasites. The results are presented as percentage (%) drug killing of parasites in at least five mice per dose.

### Probenecid and verapamil reinstate LM potency against the PQ^R^ parasites but not against the LM^R^ parasites

We screened for the effect of probenecid or verapamil on the growth of the asexual blood-stage of LM^R^ and PQ^R^ parasites (Table 2). First, we assessed the activity of PQ against the WT parasites. PQ at 6mg/kg and 1.5mg/kg killed 98.69% and 57.96%, respectively, confirming the potency of the drug (Table 3). Verapamil [50mg/kg] yielded 0% and 0.81% against the LM^R^ and PQ^R^ parasites, respectively. Probenecid [400mg/kg] yielded 8.44 % and 8.31% against LM^R^ and PQ^R^ parasites. Probenecid [400mg/kg] and verapamil [50mg/kg] were therefore selected for combination assays with either LM or PQ. We first tested the impact of probenecid or verapamil on LM potency against the LM^R^ parasites (Fig 3B). While 100mg/kg of LM yielded 23% parasite killing, a combination of probenecid [400mg/kg] and LM [100mg/kg] yielded 28.2%. Verapamil [50mg/kg] and LM [100mg/kg] yielded 26.5%. These results suggest that probenecid or verapamil failed to restore LM potency against LM^R^ parasites. Next, we measured the impact of probenecid or verapamil on PQ activity against the PQ^R^ parasites (Table 3). While PQ [75mg/kg] showed a meager 8.39% parasite killing, a combination of PQ [75mg/kg] with either probenecid [400mg/kg] or verapamil [50mg/kg] yielded 0% parasite killing (Table 3). These results show that both probenecid and verapamil did not potentiate PQ activity against the PQ^R^ parasites.

Since the PQ^R^ parasites are also highly resistant to LM^26^, we evaluated whether probenecid or verapamil could restore LM potency against PQ^R^ parasites. As expected, LM [50mg/kg] had 0% parasite killing. To our surprise, probenecid at 400mg/kg reinstated the parasite killing potency of LM [50mg/kg] from 0% to above 60% (p<0.0001) (Fig 3B). In a familiar trend with other reversing agents, 50mg/kg of verapamil significantly re-established LM [50mg/kg] potency against the PQ^R^ parasites from 0% to above 55% (p<0.0001) (Fig 3B). This selective restoration of LM by probenecid, cyproheptadine, and verapamil against the PQ^R^ parasites formed the basis for our subsequent molecular studies.

### Essential drugs transporters and enzymes are differentially expressed between drug-resistant and wildtype parasites

To dissect the mechanisms of PQ and LM resistance and reversal, we hypothesized that major transporters and enzymes involved in the transport and facilitation of drug action could mediate LM and PQ resistance. These genes include; the *crt, mdr1, mrp2, pank, fnr, pmiv, pmix, pmx, SufS, and pi3k*. In PQ^R^ parasites, the mRNA profile showed that the expression of two genes, *SufS*, and *pmix* was upregulated by 1.47-fold (p<0.018) and 2.08-fold (p<0.007), respectively (Fig 4). The mRNA expression was suppressed in nine genes in the LM^R^ parasites, save for the pmiv gene, which showed marginal elevation of the mRNA amounts (1.04fold (p<0.56) (Fig S1 https://osf.io/rfvgp). After profiling and comparing the gene expression trends between the LM^R^ and PQ^R^ parasites, we found that SufS, pmiv, and pmix had differential expression trends between the two drug-resistant parasites. Overall, we recorded a lower expression of mRNA in *crt, mdr1, mrp2, fnr, pmx*, and *pi3k* genes in both LM^R^ and PQ^R^ parasites. Two genes, pmix, and sufs were uniquely elevated and warranted further investigations. To date, pmix has not been associated with the susceptibility of current antimalarial drugs; however, genes in Fe-S biogenesis and FNR redox systems have been associated with ACTs susceptibility such as artemether and lumefantrine ^30,31^. Additionally, we recently mapped a nonsynonymous mutation in the FNR protein that we associate with LM and artemisinin (ART) susceptibility in *P. berghei*(Kiboi et al. unpublished data). The FAD containing FNR is a redox couple protein of the ferredoxin (Fd) in the FNR redox system^32^. To function, the FNR redox system relies on the supply of Fe-S clusters supplied by SufS and associated enzyme SufE within the apicoplast ^33^. In this study, we selected FNR and SufS protein for further evaluation. Since FNR was downregulated while SufS was overexpressed in the PQ^R^ parasites, we sought to reverse these mRNA expression trends and then measure the impact on drug susceptibility.

**Fig 4:**
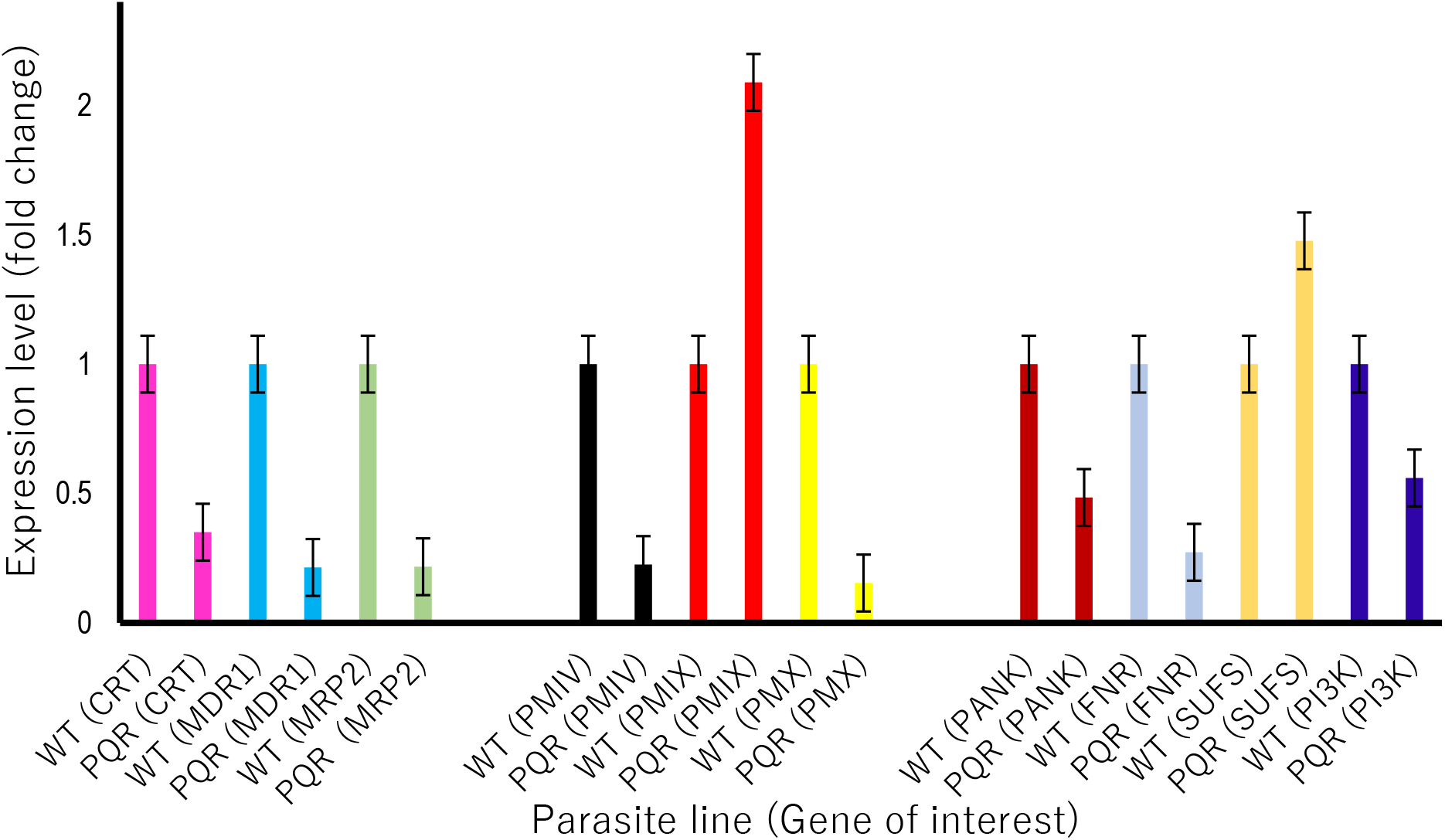
Gene expression profiles of transporters and enzymes: Transporters: chloroquine resistance transporter (CRT), multidrug resistance protein (MDR1), multidrug resistance-associated protein 2 (MRP2). Enzymes: Plasmepsin IV (PMIV), Plasmepsin IX (PMIX), Plasmepsin X (PMX) or phosphatidylinositol 3-kinase (PI3K), pantothenate kinase (PANK), ferredoxin NADP-reductase (FNR), cysteine desulfurase (SUFS), as measured from cDNA amount derived from 5*μ*g/μl of total RNA isolated from piperaquine resistant clones (PQR) relative to the wild-type piperaquine sensitive (PQS). The significance of differential gene expression from a mean of three independent experiments was computed using the Mann–Whitney U test with p-value set at 0.05.

### The successful generation of SufS knockdown or FNR over-expression transgenic parasites

We sought to investigate the impact of altering mRNA expression of FNR or SufS in the PQ^R^ parasites on LM susceptibility. We successfully generated by transfection two transgenic parasites: one with a deleted SufS gene (PQ^R^_SufS_KD) and the other with an overexpressed FNR gene (PQ^R^_FNR_OE) (Fig 5). Recovered parasites after transfection were cloned by limiting dilution to generate genetically homogenous parasites and genotyped using three sets of primers: QCR2/GW2, GT/GW1, and QCR1/QCR2. As expected, using the vector-specific QCR2 and the standard GW2 primers, we obtained a fragment of 0.9kb, and 1.5kb, respectively, in PQ^R^_SufS_KD (Fig 5A) and PQ^R^_FNR_OE parasites (Fig 5B). PCR amplification analysis using vector-specific quality control primers, the QCR1, and QCR2 amplified in both PQ^R^ parasites and PQ^R^_SufS_KD parasites (Fig 5A) as well as in both PQ^R^ and PQ^R^_FNR_OE parasites (Fig 5B) confirming that the SufS knock out vector, as highlighted from PlasmoGEM ^34^, does not cover the entire gene thus a possibility of generating a knockdown transgenic parasites. To confirm integration of the vector into the correct chromosome and position, PCR analysis using standard primers GW1 and the vector-specific primers GT yielded, as anticipated, PCR products of 1.9kb and 3.9kb for PQ^R^_SufS_KD (Fig 5A) and PQ^R^_FNR_OE parasites, respectively (Fig 5B). To confirm the overexpression of the FNR, we quantified the mRNA expression in PQ^R^_FNR_OE line relative to the PQ^R^ parasites. The results indicated a 5-fold (p<0.0046) increase in the mRNA amount of the FNR gene in the PQ^R^_FNR_OE parasites (Fig 5C). To assess the partial disruption of the SufS gene in PQ^R^_SufS_KD parasites, the mRNA expression was quantified and robustly demonstrated a reduction in the expression of the SufS mRNA (Fig 5C). Overall, these results affirmed the successful generation of the transgenic parasites. The transgenic parasites were then utilized to measure the impact of deleting SufS or overexpressing FNR genes on the drug responses relative to the PQ^R^ and wildtype parasites.

**Fig 5:**
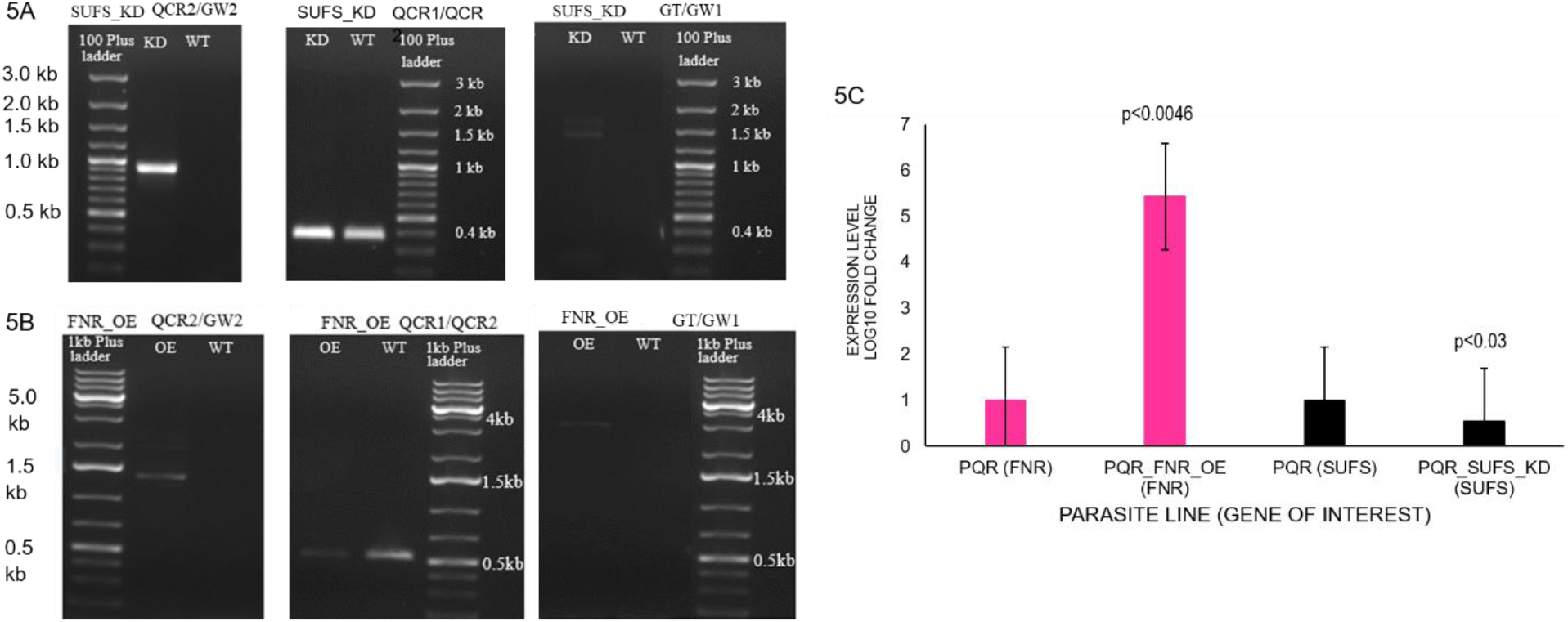
PCR genotyping of the SUFS-knockdown (KD) and FNR Over-expression (OE) parasites; **(A)** SUFS_KD, **(B)** FNR_OE and the wildtype (WT) control lines. The PCR amplification used three sets of primer pairs; QCR2/GW2 to confirm the presence of the vector, GT/GW1, to verify the integration of the vector into the parasite genome and QCR1/QCR2 to confirm the modification of the specific genes. The QCR1, QCR2, and GT are vector-specific primers, while GW2 and GW1 primers are standard primers. **(C**) Expression profiles in ferredoxin NADP-reductase (FNR), cysteine desulfurase (SUFS) as measured from cDNA amount derived from 5*μ*g/ul of total RNA isolated from transgenic parasites (PQ^R^_FNR_OE and PQ^R^_SUFS_KD) relative to the drug-resistant parent piperaquine resistant parasites (PQ^R^). The differential expression from a mean of three independent experiments was significantly different for FNR (p<0.0046), and SUFS (p<0.03) using Mann-Whitney U test with p-value set at 0.05.

### Alteration of SufS and FNR genes significantly attenuates the growth rate of the asexual blood-stage parasites

Here we hypothesized that alteration of a gene in parasites might concomitantly alter the growth rates of the asexual blood-stage parasites. To prosecute this hypothesis, we assessed the growth rates of the transgenic parasites (Fig 6). The wildtype parasites grew as expected with parasitemia reaching an average of 15% on Day four post parasite inoculation (D4 PI). Similarly, the PQ^R^ parasites had an average parasitemia of 13% on D4 PI. However, the growth rate of the transgenic parasites was significantly low. For instance, PQ^R^_FNR_OE or PQ^R^_SufS_KD yielded an average parasitemia of 8% or 10% respectively, suggesting a fitness cost on the growth of the asexual blood-stage parasites. After quantifying the growth fitness cost, the study revealed that alteration of FNR or SufS significantly reduces normal parasite growth rate phenotype by 47% and 33%, respectively.

**Fig 6:**
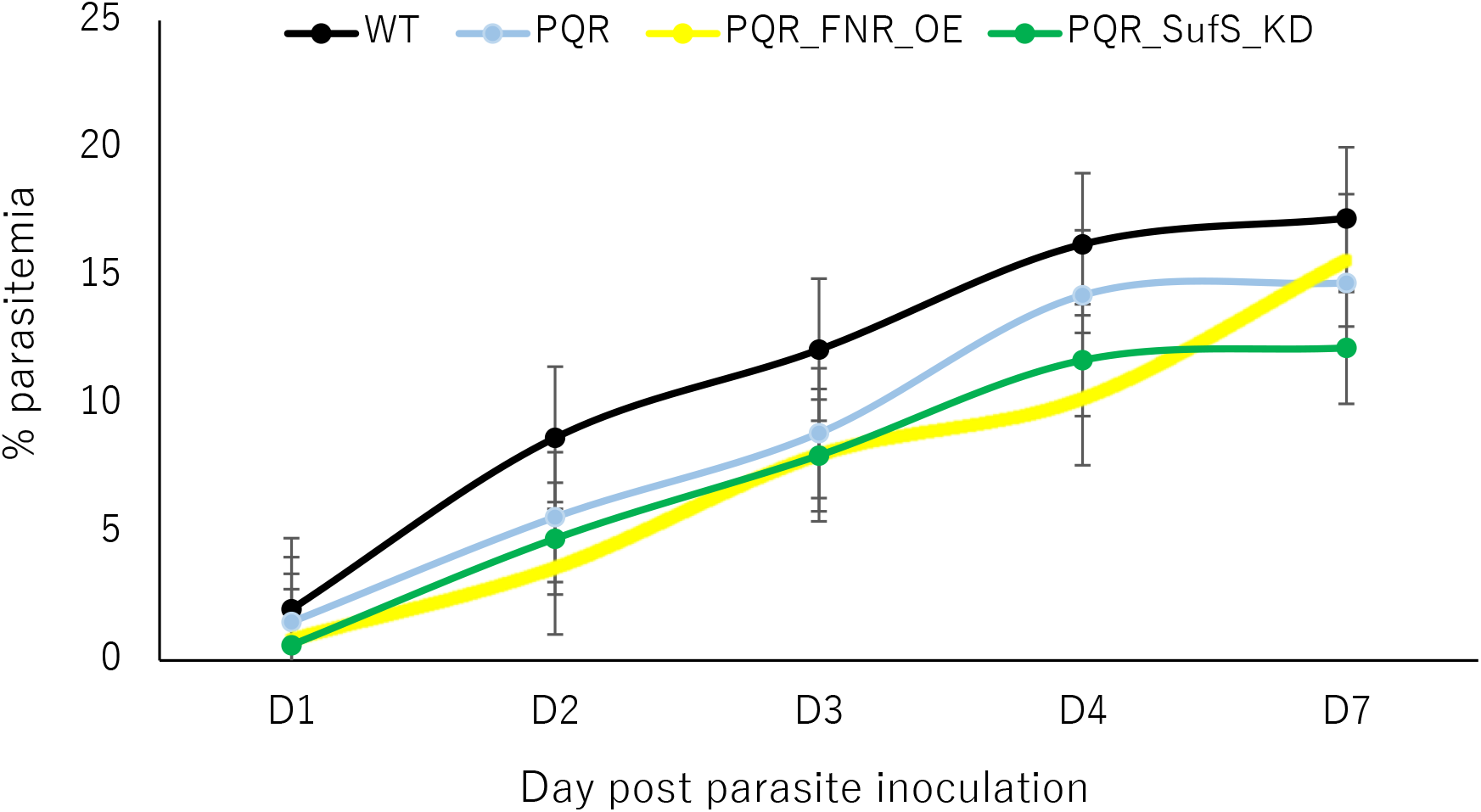
The growth rates of transgenic parasites: Average growth rate profiles of PQ^R^_SufS_KD or PQ^R^_FNR_OE relative to the wild type drug-sensitive (WT), and the piperaquine resistant (PQ^R^) parasites as determined on day 1, 2, 3, day 4, and day 7 post parasite inoculation.

### Overexpression of FNR or knockdown of SufS enhances PQ^R^ susceptibility to LM alone or in combination with the RA

To query the impact of partially deleting SufS and overexpressing FNR genes in the PQ^R^ parasites on LM and RA responses, we assessed the activity of LM alone or in combination with the RA against the transgenic and PQ^R^ parasites (Fig 7A). As expected, LM [50mg/kg] showed negligible parasite killing at 4.8% against the PQ^R^ parasites. Surprisingly, similar LM concentration regained potency against PQ^R^_FNR_OE (64.5%) and PQ^R^_SufS_KD (67.1%) parasites. Based on these results and if RA enhanced LM susceptibility via SufS and/or FNR in PQ^R^ parasites, we reasoned that the deletion of SufS and overexpressing FNR might have two biological impacts; RA might enhance LM potency against PQ^R^ parasites or the impact of RA on LM potency might be abolished. Against the PQ^R^_FNR_OE parasites, combination of LM [50mg/kg] and cyproheptadine [5mg/kg] or probenecid [200mg/kg] or verapamil [50mg/kg] yielded 62.5%, 71.8% and 53.1% respectively. In a similar trend, against the PQ^R^_SufS_KD parasites, combination of LM [50mg/kg] and cyproheptadine [5mg/kg] or probenecid [200mg/kg] or verapamil [50mg/kg] yielded 84.2%, 73.1% and 78.5% respectively. The percentage parasite killing of the RA-LM combination was not significantly different from the potency of LM alone on either PQ^R^_FNR_OE or PQ^R^_SufS_KD parasites. These results indicate that the partial deletion of SufS or overexpression of FNR in PQ^R^ parasites restored PQ^R^ parasites susceptibility to LM and abolished the action of RA on LM activity.

**Fig 7:**
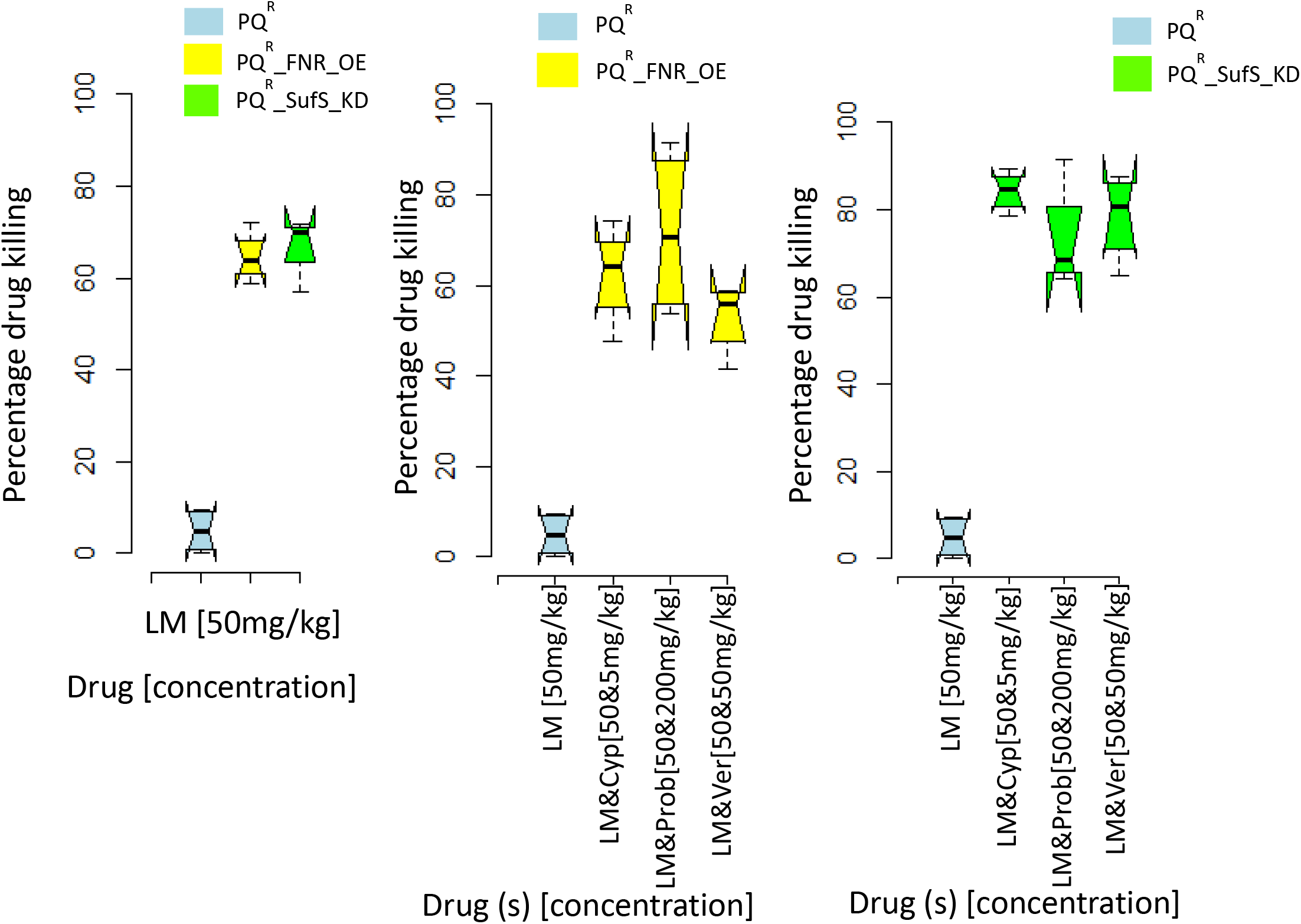
Impact of overexpressing FNR gene and deletion of SufS gene on drug response profiles as assessed in the 4-Day Suppressive Test. **(A)** The activity profile of LM alone against piperaquine-resistant parasites (PQ^R^) and transgenic parasites with an overexpressed FNR gene (PQ^R^_FNR_OE) and parasites with SufS knock down (PQ^R^_SufS_KD). **(B)** Impact of overexpressing FNR gene in the PQ^R^ on the activity of LM in combination with probenecid (Prob), verapamil (Ver), or cyproheptadine (Cyp). **(C)** Impact of knocking down the SufS gene in the PQ^R^ on the activity of LM in combination with Prob, Ver, or Cyp. The results are presented as percentage (%) drug killing of parasites from at least five mice per dose.

### Lumefantrine and cyproheptadine exhibited higher binding affinities for both FNR and SufS proteins than verapamil or probenecid

To confirm the *in vivo* drug susceptibility results and investigate the potential interaction of LM or RA with either SufS or FNR, we used *in-silico* bioinformatics tools and approaches. Here, for all the structures modeled, the Z-scores from the PROSA-web server that indicates the degree of correctness showed scores of −8.49 and −8.67 for FNR and SufS, respectively (Table 4). The results of the Z-scores obtained imply that the modeled protein structures were within the range of scores typical of the experimentally defined proteins. As expected, LM yielded high but equal binding affinities of −7.9kcal/mol on both SufS and FNR, indicating a better binding between LM and the proteins. Surprisingly, cyproheptadine showed the highest binding affinity to SufS (−9.5kcal/mol). The lowest binding affinity of −6.3 kcal/mol was between FNR and verapamil (Fig 8A and Fig 8B).

**Table 4:** Validation of the modelled 3D structures of the *Plasmodium berghei* ferredoxin NADP+ reductase (*PbFNR*) and *Plasmodium berghei* cysteine desulfurase (*PbSUFS*) proteins as determined using PROSA-web server.

| *Protein* | *Gene ID* | *Z-Score* | *score*<br>*Averaged 3D-1D score > =*<br>*0.2 (%)* | *Residues in*<br>*most*<br>*favoured*<br>*regions (%)* | *Residues in*<br>*additional*<br>*allowed*<br>*regions (%)* | *Residues in*<br>*disallowed*<br>*regions (%)* |
| --- | --- | --- | --- | --- | --- | --- |
| *PbFNR* | PBANKA_112210 | -8.49 | 86.21 | 85.9 | 13.2 | 0.4 |
| *PbSUFS* | PBANKA_061430 | -8.67 | 75.11 | 82.2 | 14.1 | 1.6 |

**Fig 8:**
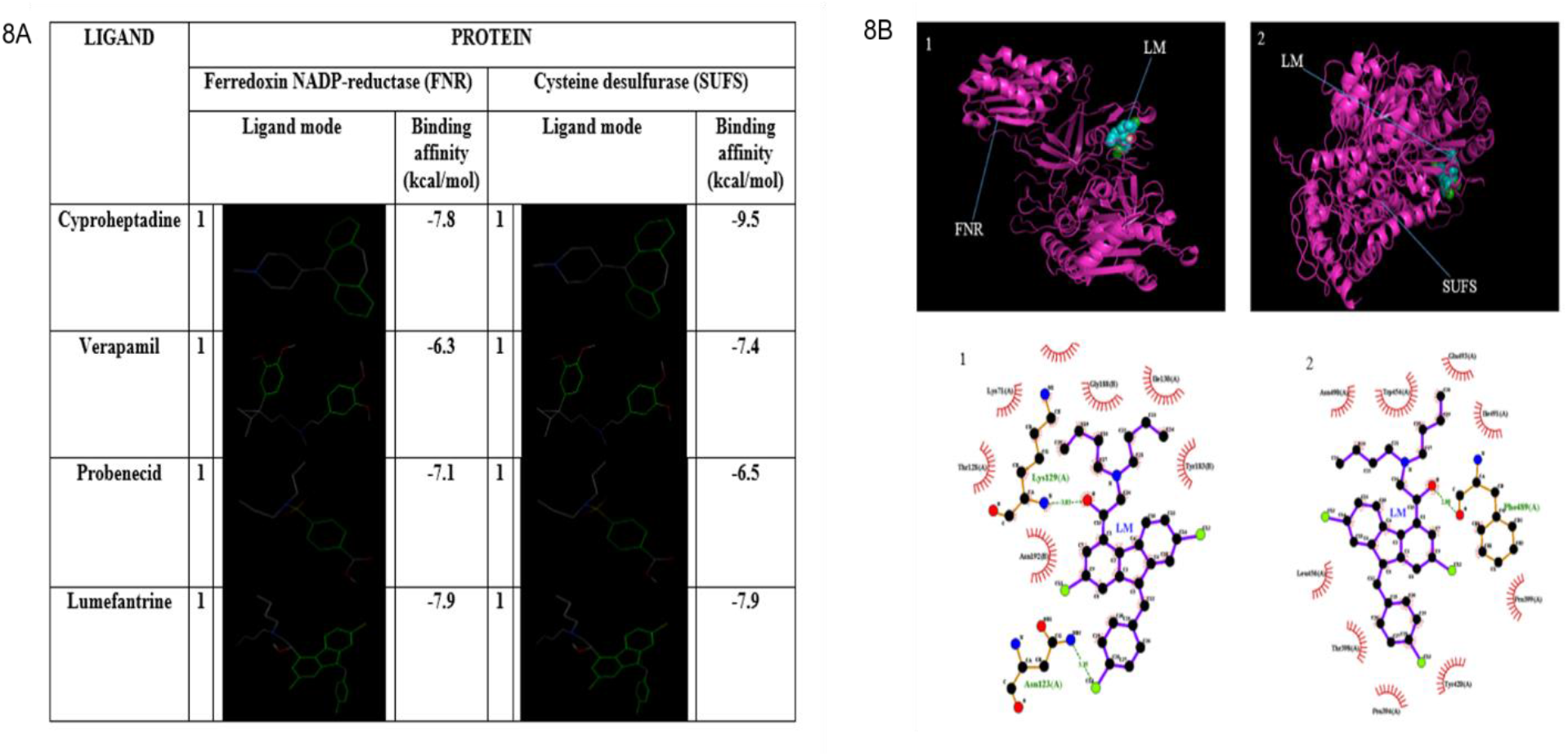
Drug binding profiles on target proteins; FNR and SUFS: **(A)** Ligand best docking modes with respective binding energies towards FNR and SUFS: LM exhibited a high binding affinity to the FNR and SUFS proteins. The binding geometries for docking of a single ligand with a single receptor as modelled using the Autodock Vina. The binding energies for LM to the FNR and SUFS were the lowest, suggesting a better affinity to the protein. Cyproheptadine yielded higher binding affinity to both FNR and SUFS as compared to the other RA, verapamil, and probenecid **(B)** Interaction of LM with the FNR and SUFS. **TOP:** The crystallographic structure of (1) FNR and (2) SUFS in complex with LM as visualised using PyMOL. **Bottom:** 2D: (1) LM-FNR and (2) LM-SUFS interaction with the oxygen atoms shown in red, nitrogen atoms shown in blue, while hydrogen bonds are shown as olive-green dotted parasites. The interaction plots were generated using LigPlot+.

## DISCUSSION

Here, we unveil new evidence that chemosensitizers: cyproheptadine, verapamil, or probenecid selectively restore LM activity against PQ^R^ parasites. We also show this selective reversal of LM resistance is abolished by the deletion of SufS and overexpression of the FNR gene. We have used parasites previously selected by submitting *P. berghei* parasites to PQ or LM for at least forty mechanical passages^35^ to systematically dissect reversal and the mechanisms of LM and PQ resistance in *P. berghei* ANKA. During the selection process, the PQ^R^ parasites were submitted to PQ pressure alone, but resultant phenotypes had significantly reduced susceptibility to CQ, AQ, LM, and DHA ^26^, displayed stability in the absence of PQ pressure and after freeze-thawing process ^28,36^. These prior studies affirmed the multidrug resistance nature of the PQ^R^ parasites. In the current study, all the chemosensitisers restored LM potency against the PQ^R^ parasites, but only cyproheptadine restored LM activity against the LM^R^ parasites. The current study, therefore, studied potential molecular mechanisms of LM resistance and subsequent potentiation of LM efficacy by the three chemosensitisers in the PQ^R^ parasites.

Our results show that cyproheptadine has at least two functions; first, and as expected, is the parasite killing ability against both the LM^R^ and the wildtype parasites, possibly by inhibition of heme polymerization as previously elaborated^15^. Second, cyproheptadine enhances parasite killing by selectively restoring LM activity against both the LM^R^ and PQ^R^ parasites. PQ and cyproheptadine exert parasite killing by inhibiting detoxification of heme in the digestive vacuole ^15^. Therefore, the low susceptibility of the PQ^R^ parasites to cyproheptadine compared to the LM^R^ and the wildtype parasites suggest that PQ and cyproheptadine also share some common resistance mechanisms. The failure of cyproheptadine to restore PQ activity against PQ^R^ parasites suggests that the PQ^R^ parasites possess at least two types of resistance mechanisms. The first mechanism codes for PQ resistance and is cyproheptadine insensitive and second, which mediates LM resistance and is cyproheptadine susceptible. It is unclear whether the genes associated with cyproheptadine modifiable mechanisms in both LM^R^ and PQ^R^ are similar or different. A lower dose of cyproheptadine [5mg/kg] had only one biological function of restoring LM activity against both LM^R^ and PQ^R^ parasites. We also note that resistance mechanisms mediating the loss of LM activity against the PQ^R^ parasites are verapamil susceptible, but the loss of LM activity against LM^R^ parasites are independent of verapamil action suggesting a complex emergence of LM resistance. Resistance to quinoline antimalarials such as PQ is associated with acidification of the digestive vacuole (DV) ^12^. Verapamil mediates reversal by replacement of positive charge and normalization of pH, blocking efflux of quinoline drugs from digestive vacuole (DV) via the mutated PfCRT protein ^37^. The mechanism of action of PQ and LM is predicted to be similar to that of standard quinoline; chloroquine ^38^. Thus, these drugs may share some resistance mechanisms. However, *PbCRT* in PQ^R^ or LM^R^ parasites were found not to carry mutations ^36^, suggesting that verapamil may use alternative proteins to enhance drug activity. Our data provide clues that verapamil may interact with FNR and SufS protein to restore and enhance LM activity in PQ^R^ parasites.

Like with verapamil and cyproheptadine, we provide compelling evidence that mechanisms of LM resistance in PQ^R^ parasites are associated with proteins that interact with organic anion inhibitors, probenecid. Probenecid is a substrate for multidrug resistance-associated protein; for instance, in *P. falciparum*, probenecid selectively sensitizes chloroquine-resistant isolates by increasing the level of chloroquine accumulation *in vitro* ^14^. Also, probenecid unselectively potentiates the activities of antifolates agents against both antifolates resistant and sensitive *P. falciparum*. This reversal of antifolates resistance is associated with inhibition of multi-resistance-associated proteins ^39^ and inhibition of endogenous folate transport ^14^. Multi-resistance associated proteins are also implicated in resistance to quinoline antimalarial drugs in both *P. falciparum* and *P. berghei* ^40^. We have previously confirmed PQ^R^ parasites as multidrug-resistant parasites; thus, probenecid mediated inhibition of multi-resistance proteins in re-establishing LM activity against PQ^R^ parasites is a possible alternative mechanism.

The next generation of antimalarial drugs has envisioned the incorporation of three drugs comprising of two long-acting drugs and a short-acting artemisinin drug ^41^. Currently, cyproheptadine, an H-1 antagonist, at 4mg/kg, is recommended for the treatment of allergic reactions and improvement of appetite (www.medscape.com). We have shown that 2.5mg/kg of cyproheptadine is a potent modulator of lumefantrine resistance in both PQ^R^ and LM^R^ parasites, at least in *P. berghei*. In humans, the recommended dosage for verapamil and probenecid is 80mg/kg and 500mg/kg, respectively (www.medscape.com). Studies have supported a combination of probenecid and SP for intermittent prevention of malaria in pregnancy (IPTp) ^13^. Here, we have used a lower dose to restore the activity of LM against PQ^R^ parasites. If the mechanisms of LM resistance in *P. berghei* and *P. falciparum* are similar, then our data demonstrate that incorporation of cyproheptadine into antimalarial combination therapy that comprises LM might achieve two aims: First is to boost LM activity and possibly delay the emergence of LM resistance. Second, an optimized cyproheptadine concentration might offer an added advantage in the combination by exerting extra antimalarial. The data also confirms the existence of more than one control channels for PQ resistance in PQ^R^ parasites. The first is associated with LM resistance and is verapamil, probenecid, or cyproheptadine inhibitable channel only in the presence of LM. The other control is linked with PQ resistance, which is verapamil, probenecid or cyproheptadine insensitive mechanisms. Currently, LM and PQ are used in ACTs therapies that are mainstay drugs for the treatment of malaria in many African countries ^42^. Again, if mechanisms of resistance to LM in *P. berghei* are the same as those in *P. falciparum*, then the emergence of LM resistance may proceed through two possible mechanisms; first is through the indirect PQ selection pressure via the PQ/DHA combination. The second mechanisms are by the direct LM pressure via the LM/ATM combination. The selective potentiation of LM potency by probenecid, verapamil, and cyproheptadine against the PQ^R^ parasites is appealing since parasites are multidrug resistant parasites and might expose shared resistance mechanisms^26^. Our data suggest that the incorporation of any of the reversing agent into an antimalarial combination therapy that comprises LM as one of the components might concomitantly antagonize the emergence of LM resistance derived from PQ selection pressure.

The impact of chemosensitizers on LM activity against both PQ^R^ and LM^R^ parasites suggests that LM resistance is under the control of more than one molecular signature and pathway. Our data show that changes in mRNA expression of SufS, a unique protein in the Fe-S biogenesis ^30^, and FNR in the FNR redox system; ^43^ are associated with PQ and LM resistance. The fact that an increase of the mRNA amount of FNR correlated with increased susceptibility of PQ^R^ parasites to LM and concomitantly abolished the impact of RA on LM activity confirm the interaction between RA and LM with FNR. Indeed, our *in-silico* binding results show significant binding power between the RA, LM, and FNR. Several studies have established that the Fd/FNR redox system, which supplies reducing power for essential biosynthetic pathways in the malaria parasite apicoplast ^44^, is a potential target for the development of specific new antimalarial agents ^43^. Validation of the FNR redox system as a potential target and resistance marker for the artemisinins was revealed by the nonsynonymous background mutation (Asp193Tyr) in the *Fd* protein in parasite exposed to the ACTs^31^. We provide a clear link between LM activity, RA action, and overexpression of the FNR gene. Redox enzymes such as FNR convert inactive compounds into active molecules ^45^,^46^. Two possible assumptions exist; First, FNR may be a target for LM. Reduced expression of FNR in the PQ^R^ parasites may potentially result in reduced binding targets for LM. The high FNR expression in the transgenic parasites PQ^R^_FNR_OE seems to restore the concentration of the FNR, thus more binding targets for LM and a consequent high susceptibility of PQ^R^_FNR_OE to LM. Second, the FNR is a potential activator for LM, therefore reduced expression of FNR in the PQ^R^ is accompanied by reduced active metabolites from LM, thus less activity. The increase in the FNR, as confirmed in the PQ^R^_FNR_OE transgenic parasites, seems to reinstate the activator; therefore, more of the LM metabolite is generated restoring LM activity in the PQ^R^_FNR_OE transgenic parasites as compared to PQ^R^ parasites. The molecular mechanism of drug-target interaction between the RA and LM with FNR requires further in-depth investigation. The reversing agents restore the activity of antimalarial drugs by enhancing the accumulation of drugs within the target site ^18^. It is likely that in the event the FNR amount is low, the threshold concentration for LM action is not achieved, the threshold may be enhanced by a RA possibly by increasing the uptake of the few active LM metabolites.

The FNR redox system relies on the Fe-S clusters for its action ^47^. Both the FNR redox system and Fe-S cluster biogenesis are essential pathways for parasite survival within the red blood cells ^33,43^, the site of action of LM and PQ. Plasmodium species possess two Fe-S cluster biogenesis pathways: the mitochondria localized ISC pathway and the SUF pathway within the apicoplast. Here, we focused on apicoplast exclusive SufS enzyme ^48^. SufS converts L-cysteine to L-alanine and sulfane sulfur via the formation of an enzyme-bound persulfide intermediate within the apicoplast compartment^49^. Here, we first confirm a significant attenuation of the growth rates on the deletion of the SufS gene, thus essential for the growth of asexual blood-stage parasites. Second, the results presented here show that an increase in the RNA amount of SufS correlates with decreased LM susceptibility in PQ^R^ parasites.

We have validated these findings through the deletion of the SufS gene in PQ^R^ parasites yielding parasite lines: PQ^R^_SufS_KD that showed significant susceptibility to LM and simultaneously obliterated the impact of cyproheptadine, verapamil, and probenecid. Taken together, our results suggest that LM interacts with SufS and exert its antimalarial action through the protein. However, this mechanism seems to be solely in the PQ^R^ parasites. We reveal clues that SufS may be a target for LM. Thus, the low expression of SufS in the LM^R^ parasites may be adaptive mechanisms for the parasites to reduce potential binding targets for the LM. Indeed, the adaptive parasite mechanism is supported by the low growth profiles of the parasite observed in this study after partial deletion of the SufS in the PQ^R^ parasites. Abolishing of the impact of chemosensitiser on LM activity, expose additional route for reversal of resistance using chemosensitisers. Our results on mRNA expression of SufS seemed antagonistic between PQ^R^ and LM^R^ parasites. These findings reveal that LM resistance may associate with SufS if the selection pressure applied is PQ; however, LM pressure does not select high expression of SufS, and thus resistance to LM through LM pressure is independent of SufS protein.

In conclusion, this study provides critical evidence regarding LM resistance in a PQ resistant *P. berghei* ANKA; As highlighted earlier, PQ pressure yields multidrug resistant PQ^R^ parasites that are accompanied by the loss of LM efficacy ^26,28^. First, we have restored LM efficacy against the PQ^R^ parasites by combining with either cyproheptadine, verapamil, or probenecid. SufS deletion and FNR overexpression in the PQ^R^ parasites abolishes the impact of the chemosensitizers. We provide essential clues and propose the association of SufS and FNR with the action of LM and chemosensitizers. Taken together, if mechanisms of LM resistance between *P. berghei* and *P. falciparum* are similar, then the mRNA expression of SufS and FNR genes may be predictors of the emergence of LM resistance in the *P. falciparum* field isolates. Also, our data suggest that the incorporation of any of the chemosensitizers into an antimalarial combination that comprises LM might augment LM activity and concurrently antagonize the emergence of LM resistance derived from PQ pressure. The in-depth mechanisms of how SufS and FNR mediate LM resistance and the impact of their differential expression in drug susceptibility should be investigated in the human malaria parasite *P. falciparum*.

## MATERIALS AND METHODS

### Parasites, Host, and Compounds

We used two parasite reference lines *of P. berghei* ANKA, the MRA-865 and MRA-868 reference lines obtained from MR4, ATCC^®^ Manassas, Virginia, which were used as the drug-susceptible wildtype (WT) parasites. Also, two multidrug-resistant parasite lines; the PQ-resistant (PQ^R^) and LM-resistant (LM^R^) *P. berghei* ANKA parasites previously selected from MRA-865 and MRA-868 lines, respectively^26^ were studied. The two parasite lines were confirmed as stable resistant parasites^26,28^. Male Swiss albino mice weighing 20±2g outbred at KEMRI, Animal house Nairobi were utilized, housed, and fed as previously described^50^. The knockout (KO) vector (PbGEM-018972), as well as the overexpression (OE) vector (PbGEM-456502), were kindly provided by the *PlasmoGEM* project under an agreement of a material transfer (PG-MTA-0093). Graphical overview of KO and OE vector design are outlined in PlasmoGEM (https://plasmogem.sanger.ac.uk/), and experimental strategies have previously been detailed^34^. Verapamil, probenecid, and cyproheptadine were purchased from Carramore international limited (UK). LM and PQ were donated from Universal Corporation, Kikuyu, Kenya. All drugs, PQ, LM, verapamil, probenecid, and cyproheptadine, were freshly prepared and administered as previously described^36^.

### Ethical consideration

This study was conducted at KEMRI and JKUAT. All mouse experiments were carried out as per relevant national and international standards (The ARRIVE Guidelines) and as approved by KEMRI-Animal Use and Care Committee. This study was permitted and ethically approved by KEMRI under the SERU No 3914.

### Drug sensitivity assays

The activity profiles of probenecid, verapamil, or cyproheptadine alone and in combination with PQ, or LM against the wildtype, resistant, or the transgenic parasites were assessed in the standard 4-day suppressive test (4-DT) in which the parasite is exposed to four daily drug doses^51^. Infection, randomization of experimental mice, drug administration, estimation of parasite densities, and calculation of percentage drug killing was performed and determined as previously described^26,50,52^.

### Extraction of RNA, synthesis of cDNA, and qPCR assays

The quantification of mRNA transcripts of *Pbcrt, Pbmdr1, Pbmrp2, Pbpank, Pbfnr, PbSufS, Pbpmiv, Pbpmix, Pbpmx*, and *Pbpi3k* was carried out using primer listed on Table S1 https://osf.io/j5w3e. Extraction of total RNA from the wildtype, LM^R^, PQ^R,^ and transgenic parasites and synthesis of respective first-strand cDNA was performed as previously described^50^. Using the *Pbβ-actin I*, as the housekeeping gene, the relative expression profile of the ten genes was performed in a final volume of 20*μ*l using Maxima SYBR Green/ROX qPCR Master Mix (Thermo Scientific™). Briefly, 12μl of Maxima SYBR mix, 3μl water were mixed with 2.0μl (0.25μM) of forward and reverse primers each, and 1μl cDNA. The reaction mix was run for pre-treatment at 50°C, for 2 min; initial denaturation at 95°C for 10 min; denaturation at 95°C for 15 secs; and annealing at 60°C for 60 secs for 45 cycles.

### Generation of the transgenic parasites

#### Isolation, digestion, and purification of the vector DNA

Using the QIAfilter Plasmid Midi Kit (Qiagen™), Plasmid DNA for each of the vectors was isolated from cultures of *E. coli* (vector hosts) in terrific broth (TB) supplemented with kanamycin after overnight culture. Following a standard protocol, the extracted plasmid DNA was restricted using Not I enzyme to release the vector backbone, followed by concentration using conventional ethanol precipitation protocol^53^. The vector was dried for 5 min at 65°C and then dissolved in 10*μ*l of PCR water. Besides Not I restriction digest, a diagnostic PCR was performed using standard primer pair (GW2 and a vector-specific QCR2)^34^ to verify the isolated vector of interest.

#### Purification of the *P. berghei* schizonts for transfection

Using standard *in vitro* culture protocols as described by ^54^, we collected *P. berghei* schizonts for transfection. Briefly, at least three mice were used for schizont culture for each of the vectors. Propagation of *P. berghei* parasites was done by intraperitoneal (IP) injection into the mice, and parasites were harvested using cardiac puncture for schizont culture at 3% parasitemia. For schizonts propagation and subsequent purification, rings stage parasites (the asexual blood-stages) were incubated at 37°C in vitro as reported previously^52,55^. Harvested and purified schizonts were re-suspended in schizont culture media, ready for transfection.

#### Transfection and selection of transgenic parasite lines

A SufS specific knockout vector (PbGEM-018972) and an FNR overexpression vector (PbGEM-456502) were separately transfected in the PQ^r^ parasites. Each of the transfection mixes contained, 20*μ*l of schizonts, 10*μ*l (5*μ*g) of plasmid DNA, and 100ml of supplemented nucleofector solution (Amaxa™). Exactly 100*μ*l of the transfection mix in Amaxa cuvette was electroporated in a Nucleofector 2B Device (Lonza™) using the program U33. A naïve mouse was intravenously injected with the electroplated parasites. Three transfections were performed for each of the vectors to increase the chance of recovering transgenic parasites. Pyrimethamine drug at 7 mg/mL, for the selection of transgenic parasites, was provided in drinking water, 24hours post-parasite inoculation for a total of 12 days. Freshly prepared pyrimethamine drug was provided every 72 hours. After the recovery of transgenic parasites, genetically homogenous parasite lines were generated by dilution cloning protocol ^36,54^.

#### Genotyping of transgenic parasites

Genomic DNA (gDNA) was extracted from the transgenic parasite lines using the QIAamp DNA Blood Mini Kit (Qiagen™) following manufacturer instructions. Three pairs of primers were used for diagnosis and genotyping, as detailed in Table S2 https://osf.io/zkxyj:i) QCR2/GW2 pair ii) QCR1/QCR2 and iii) GT/GW1 or GW2. Each reaction mixes contained 1μl of gDNA, 12.5μl of 2xGoTag Green master mix (Promega™), 6.5μl of water and 2.5μl of each of the primers. The PCR reactions were run using a 2xGoTag Green master mix for thirty cycles with an annealing temperature of 50° C and 62° C as the extension temperature. The PCR products were resolved on a 1% agarose gel.

### Molecular docking

#### Homology modeling

The 3D structures of the SufS; PBANKA_0614300, and FNR; PBANKA_1122100, were predicted by selecting as templates, the best structures with the highest similarity to their sequences available in NCBI protein database http://ncbi.nlm.nih.gov and *Plasmodium* database http://plasmodb.org/. SWISS-MODEL ^56^ available at https://swissmodel.expasy.org/ was used to predict the homology models. The SWISS-MODEL was a preferred modeling server because it annotates essential cofactors and ligands as well as quaternary structures, allowing for modeling of complete structures with less complicated software packages ^56^. The structures were then downloaded and saved in PDB format.

#### Structure validation

The models were analyzed using PROCHECK ^57^ and PRoSA-web ^58^ to determine the quality of the modeled structures. PROCHECK was used to evaluate the general stereochemistry of the protein, while PRoSA-web was employed to assess for potential errors in the 3D structures. Verify_3D was used to determine the compatibility of an atomic model (3D) with its amino acid sequence (1D)^59^.

#### Ligand selection

To perform docking experiments, the most potent RA, as well as LM, were selected as test ligands. The molecular structures of the four chosen ligands were downloaded from the ChemSpider (http://www.chemspider.com/), an online database to access unique chemical compounds ^60^. For compatibility with the docking software (https://cactus.nci.nih.gov/translate/), we converted the chemical structures from the Mol2 format into PDB format using the CADD Group’s Chemoinformatics Tools and User Services

#### Binding site analysis and ligand docking

PyMOL version 1.6.x (Delano, 2002) was used to visualize the modeled structures and binding sites. The grid box parameters of the ligands and the receptors optimization were executed using scripts within AutoDock Tools (ADT). The files were then saved in PDBQT format, and their corresponding coordinates rewrote into a configuration file used for docking. The configuration file specified the pdbqt files for both the ligand and the proteins as well as the docking parameters (dimensions and spacing angstrom). Optimization of the inputs involved the removal of water molecules from the receptor and the addition of missing hydrogen ions. Docking of a single receptor and a single ligand was carried out using the Autodock Vina ^61,62^. Autodock Vina employs the Lamarckian genetic algorithm (LGA). FNR and SufS, whose 3D structures were available in the protein data bank (PDB; http://www.rcsb.org/pdb/explore.do?structureId=2hvp) were also included. The binding energies and the positional root-mean-square deviation (RMSD) of the proteins and the ligands were presented on an output file. A ligand orientation with low binding energy signifies a better affinity towards a receptor.

### Data presentation and statistical analysis

The percentage parasitemia and the percentage of drug killing were analyzed using the Nonparametric Mann-Whitney U Test in the R statistical software with a *p*-value set at 0.05. The means for the gene expression levels from three independent experiments and triplicate assays were compared using the Mann-Whitney U test with a p-value set at 0.05. The relative expression level data were normalized using *β*-actin I as the housekeeping gene based on the formula 2^ΔΔCT63^.

## Data Availability

All lumefantrine reversal datasheet files are available from the https://osf.io/brz63/

All piperaquine reversal datasheet files are available from the https://osf.io/5trab/

All PQR_FNR_OE parasites datasheet files are available from the https://osf.io/pdvj6/

All PQr_SufS_KD parasites datasheet files are available from the https://osf.io/8upkc/

## Acknowledgments

We thank the Wellcome Trust Sanger Institute *PlasmoGEM* project team for providing the highly efficient gene modification resources used in this study.

---

**Fig S1:**
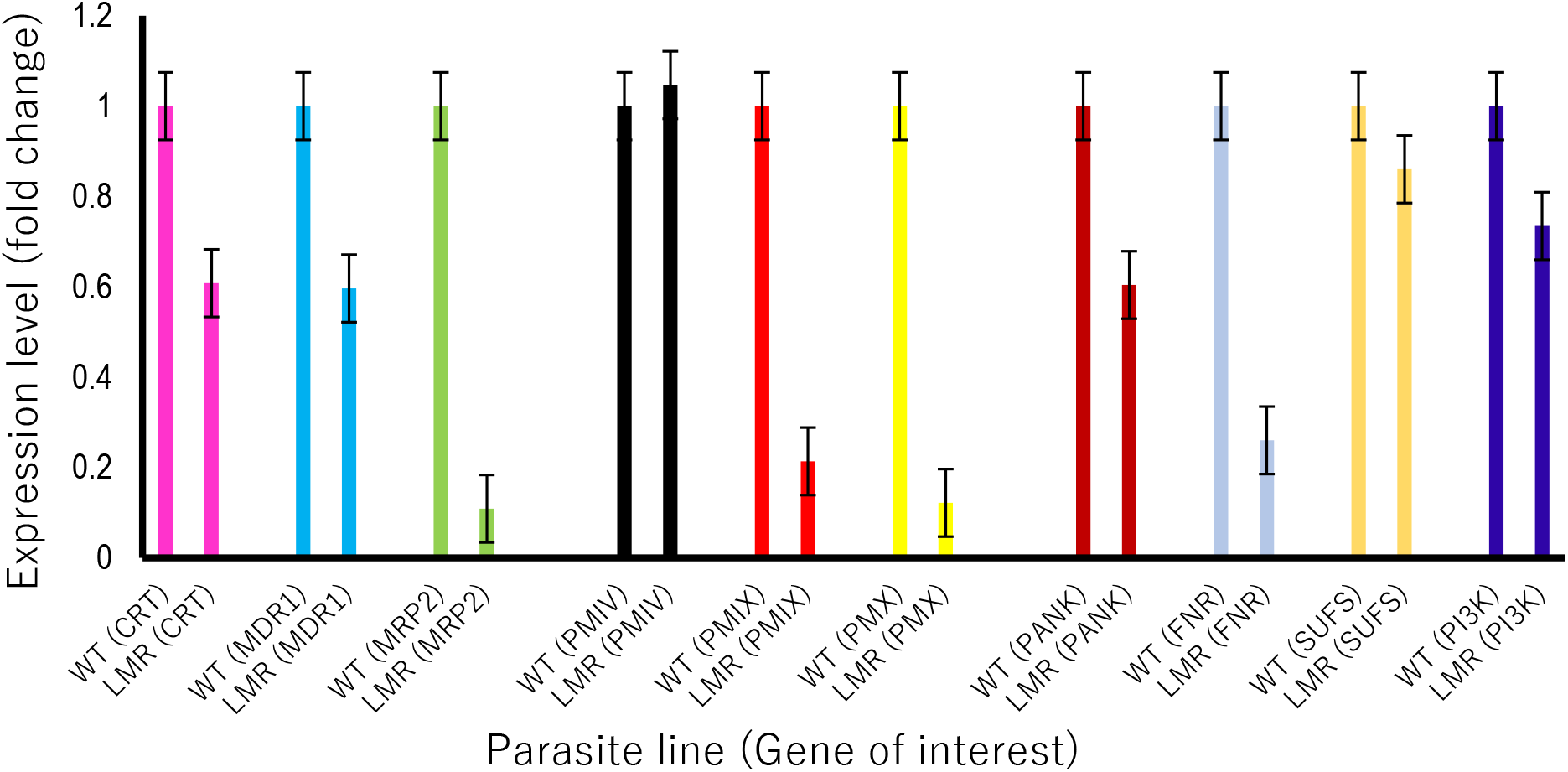
Gene expression profiles of transporters and enzymes in lumefantrine resistant Plasmodium berghei ANKA: Transporters: chloroquine resistance transporter (CRT), multidrug resistance protein (MDR1), multidrug resistance-associated protein 1 (MRP2). Enzymes: Plasmepsin IV (PMIV), Plasmepsin IX (PMIX), Plasmepsin X (PMX) or phosphatidylinositol 3-kinase (PI3K), pantothenate kinase (PANK), ferredoxin NADP-reductase (FNR), cysteine desulfurase (SUFS), as measured from cDNA amount derived from 5*μ*g/μl of total RNA isolated from lumefantrine resistant clones (LMR) relative to the wild type lumefantrine sensitive (LMS). The differential expression from a mean of three independent experiments were significantly different for CRT (p<0.0013), MDR1 (p<0.039), MRP2 (p<0.001), FNR (p<0.0026), PM9 (p<0.023), PM10 (p<0.008), PANK (p<0.046), and PI3K (p<0.013) but not significantly different for SUFS (p<0.095), PM4 (p<0.56) using Mann–Whitney U test with p-value set at 0.05.

**Table S1:**
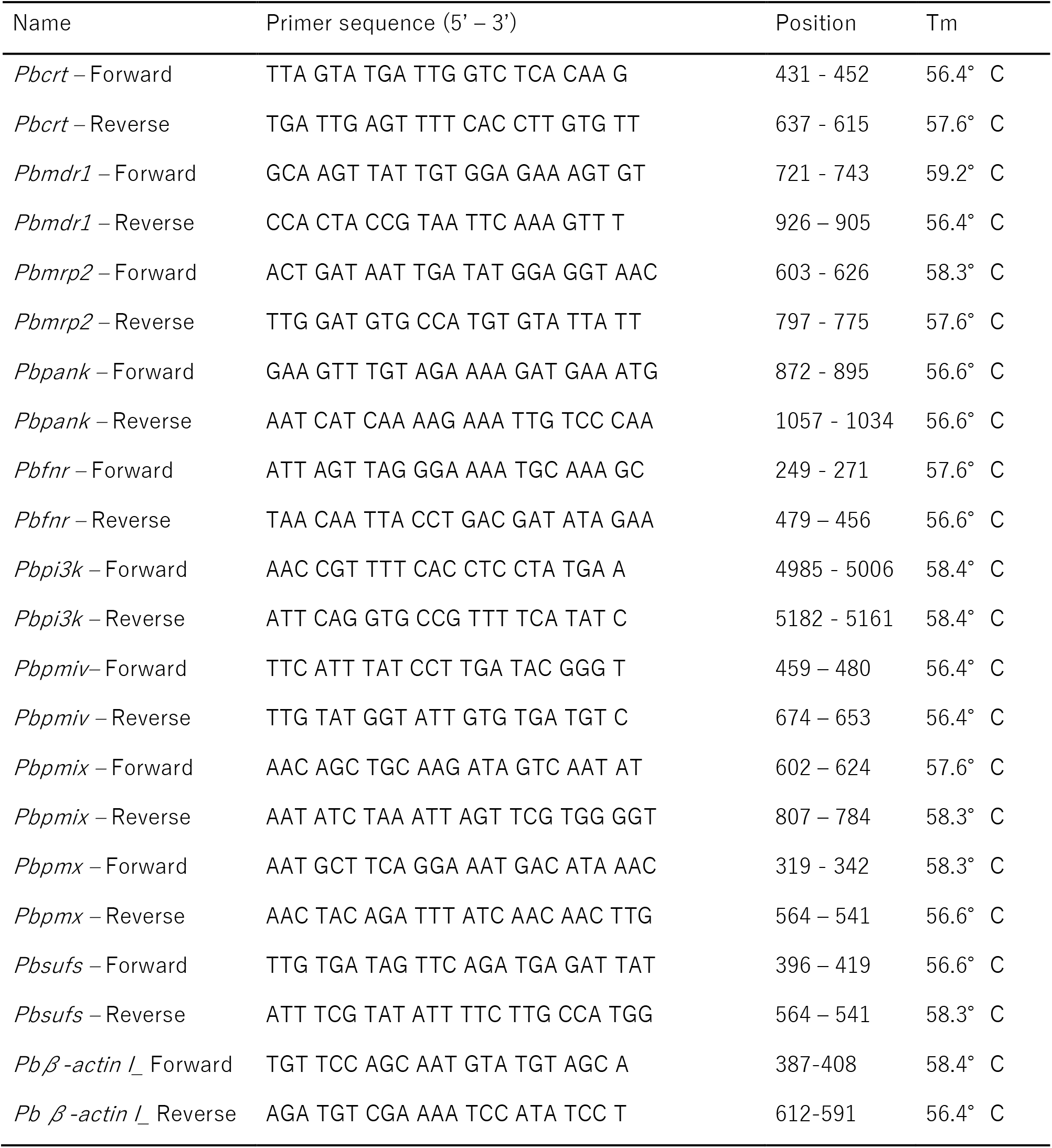
Oligonucleotide primers sequences used to measure the transcriptional level profiles of chloroquine resistance transporter (*crt*) multidrug-resistant 1 (*mdr1*), multidrug resistance-associated protein 2 (*mrp2*), pantothenate kinase (*pank*), ferredoxin NADP^+^-reductase (*fnr*), cysteine desulfurase (*sufs*), Plasmepsin IV (*pmiv*), Plasmepsin IX (*pmix*), Plasmepsin X (*pmx*) or phosphatidylinositol 3-kinase (*pi3k*) with *β*-actin I as the housekeeping gene using Maxima SYBR Green chemistry in qPCR.

**Table S2:**
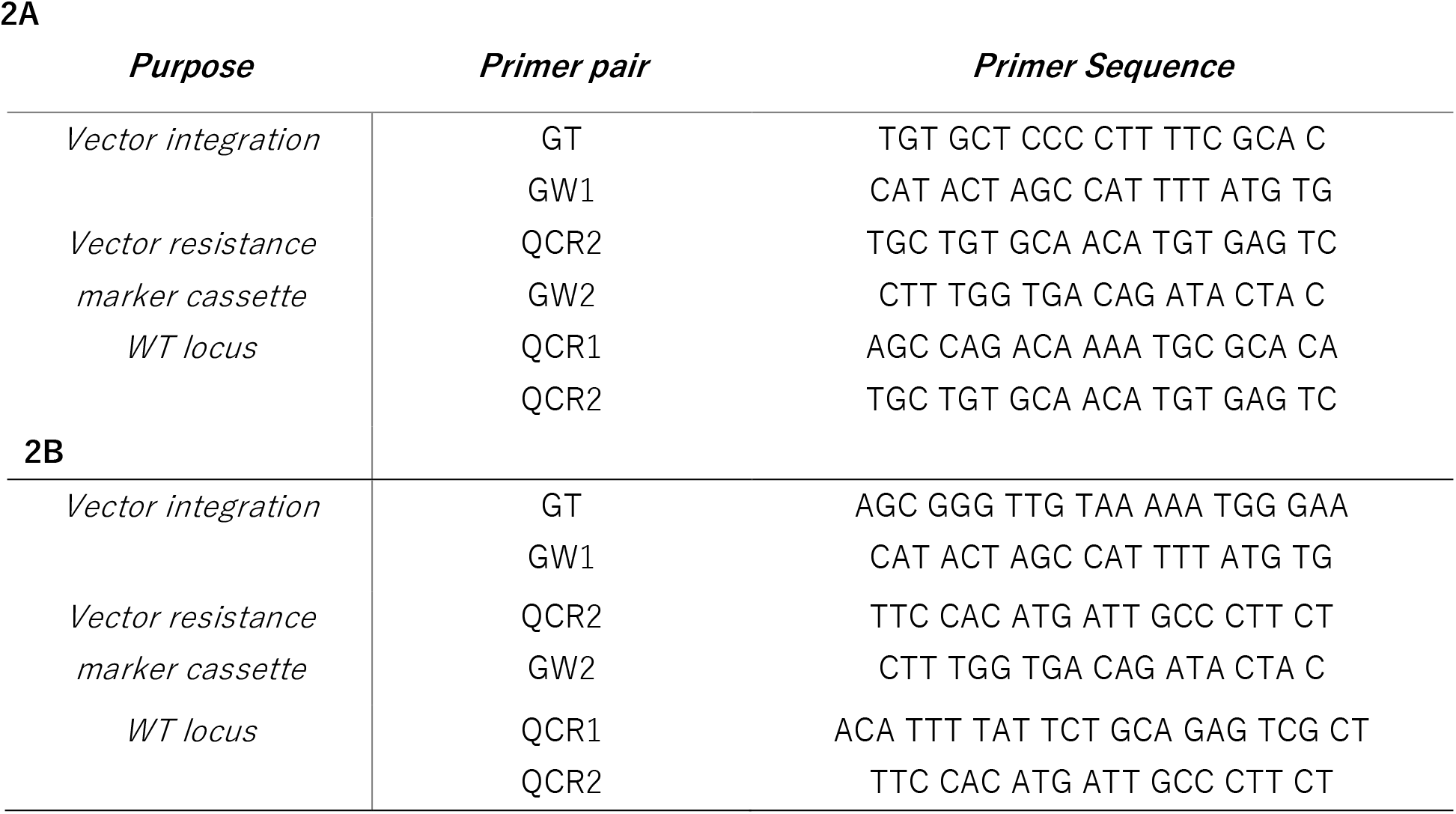
Oligonucleotide primer sequences used for genotyping of the transgenic parasite lines. (A) PQ^R^_SUFS_KD transgenic parasites. (B) PQ^R^_FNR_OE transgenic parasites.

